# Trisomy 21 alters ciliary localization of Sonic Hedgehog signaling proteins

**DOI:** 10.64898/2026.02.06.704423

**Authors:** Wolfgang E. Schleicher, Bailey L. McCurdy, Cayla E. Jewett, Chad G. Pearson

## Abstract

Trisomy 21 (T21), the cause of Down syndrome, impairs cilia-dependent Sonic Hedgehog (SHH) signaling. We investigated how T21 affects SHH signaling at the primary cilium and found that Smoothened (SMO) protein - the main effector of SHH signaling - is diminished in T21 primary cilia. Specifically, T21 cells exhibit both decreased SMO localization to cilia and reduced ciliary SMO phosphorylation (pSMO). Our findings suggest that reduced ciliary SMO in T21 results from defective entry into or maintenance in the cilium rather than impaired transport to centrosomes. Moreover, unlike the ciliogenesis defects observed in T21, the SMO defect does not appear to be associated with increased PCNT protein abundance. We also found that GPR161 (a negative SHH regulator), exhibits increased ciliary localization in T21. Notably, prolonged serum depletion to promote primary cilia maturation equalizes ciliary levels of SMO and GPR161 between T21 cells and D21 controls. We propose that delayed primary cilia maturation contributes to defective SHH signaling in T21.

**SUMMARY:** T21 impedes primary cilia assembly and disrupts SHH signaling by diminishing ciliary SMO and increasing GPR161, but extended ciliary maturation equalizes SHH protein localization between D21 and T21.

## INTRODUCTION

Down syndrome is the most common chromosomal abnormality, affecting an estimated 1 in 1000 childbirths (Weijerman and de Winter, 2010). It is caused by trisomy of human chromosome 21 (Hsa21; T21) producing phenotypes such as craniofacial abnormalities (Richtsmeier et al., 2000), cerebellar hypoplasia (Haydar and Reeves, 2012), intellectual disability (Haydar and Reeves, 2012; Antonarakis et al., 2020), and congenital heart defects (Bergström et al., 2016). Many of these phenotypes overlap with processes regulated by Sonic Hedgehog (SHH) signaling which drives organogenesis, and craniofacial development (Rowton et al., 2022; Park et al., 2019; Abramyan, 2019; Jing et al., 2023). Depending on the context, SHH promotes cell proliferation or differentiation (Zhang and Beachy, 2023; Irigoín and Badano, 2011; Rowton et al., 2022). While there are no known SHH signaling components encoded on Hsa21, individual overexpression of 163 genes located on chromosome 21 revealed that some chromosome 21 resident genes enhance SHH signaling while others suppress it (Moyer et al., 2023). This reflects the genetic complexity of both Down syndrome and SHH signaling.

Understanding changes to SHH signaling in T21 is complicated by tissue and cell specific SHH responses, which can either trigger proliferation or differentiation in target cells. T21-associated developmental defects in the brain demonstrate these context specific SHH disruptions. T21 diminishes the proliferative response to SHH in murine granule cells, resulting in reduced cerebellar size and complexity (Roper et al., 2006; Jewett et al., 2023). T21 also diminishes the morphogenic response to SHH in human neural progenitor cells, resulting in impeded oligodendrocyte differentiation (Klein et al., 2021). Although genetic and epigenetic effects of T21 help explain the clinical features of Down syndrome (Bull Marilyn J., 2020), its interactions with SHH signaling are complex and multifaceted. Therefore, understanding the cell biology of T21 is critical to decipher alterations to SHH signaling and the subsequent consequences for proliferation and differentiation that drive development.

SHH signaling relies on coordinated localization of SHH pathway proteins within primary cilia. Primary cilia extend from the cell surface with a microtubule-based axoneme ensheathed by a specialized membrane composed of unique proteins and phospholipids that facilitate ciliary signaling. In the absence of SHH ligand, the SHH receptor, Patched (PTCH) localizes to cilia and prevents ciliary accumulation and activation of the G-protein coupled receptor (GPCR) and SHH effector, Smoothened (SMO; (Rohatgi et al., 2007)). Furthermore, ubiquitination of SMO can promote its proteasomal degradation or direct its localization, ensuring transience within the cilium (Desai et al., 2020; Shinde et al., 2020; Lv et al., 2021, 2022). SHH signaling is additionally suppressed by ciliary GPR161. GPR161 is a constitutively active GPCR that contributes to a pool of cAMP utilized by PKA-C to phosphorylate Gli2/3, thus promoting its proteolytic processing into a transcriptionally repressive form (GliR; (Mukhopadhyay et al., 2013; Wang et al., 1999, 2000; Pan et al., 2009)).

In contrast, when SHH signaling is activated, signal transduction is initiated by the binding of SHH ligand to PTCH (Rohatgi et al., 2007). Upon binding SHH ligand, PTCH exits the primary cilium, allowing for deubiquitination, ciliary accumulation, and activation of SMO (Milenkovic et al., 2009, 2015; Corbit et al., 2005; Bangs and Anderson, 2017; Rohatgi et al., 2007). SMO activation depends on two events: cholesterol binding at the extracellular cysteine-rich domain of SMO that induces its protein conformational change (Kinnebrew et al., 2022; Huang et al., 2016), and phosphorylation of SMO’s C-terminal tail by G protein-coupled receptor kinase 2 (GRK2; (Walker et al., 2024)). When activated, SMO binds and sequesters PKA-C, preventing Gli2/3 phosphorylation and subsequent proteolytic processing into GliR (Arveseth et al., 2021; Happ et al., 2022; Walker et al., 2024; Niewiadomski et al., 2014). The interaction between SMO and PKA-C is maintained as SMO and Gli2/3 are transported along the ciliary axoneme by anterograde intraflagellar transport (IFT). At the distal end of the cilium, Gli2/3 is dissociated from the inhibitory partner, Suppressor of Fused (SUFU), yielding a transcriptionally active Gli complex (GliA). GliA is transported to the cell body via retrograde IFT and localizes to the nucleus to transcribe SHH target genes (Wheway et al., 2018; Goetz and Anderson, 2010).

Given that SHH signaling relies on localization and activity of several proteins within the cilium, successful construction of the primary cilium is critical. T21 disrupts primary ciliogenesis due to mislocalization of ciliary proteins caused by the overexpression of the human chromosome-21-resident gene, Pericentrin (PCNT; Galati et al., 2018; Jewett et al., 2023; McCurdy et al., 2022). Reduced ciliary SMO has been observed in mouse models of Down syndrome and patient-derived T21 fibroblasts (Jewett et al., 2023; Galati et al., 2018). However, the mechanism of how T21 reduces ciliary SMO, and whether elevated PCNT directly affects SHH signaling, remains unknown.

In this study, isogenic human hTERT-RPE-1 cell lines were employed to investigate how SHH signaling is altered in T21. SMO phosphorylation and localization to the primary cilium is diminished in T21. Moreover, T21 elevates ciliary levels of GPR161 and diminishes its ciliary exit upon SHH signal activation. This diminished localization of SMO to the cilium in T21 cells is not due to a defect in SMO transport to the centrosome and is independent of increased PCNT. Finally, ciliary SMO levels and ciliary GPR161 exit can be normalized with increased time in serum-depleted medium, suggesting that T21 primary cilia are delayed in maturation and effective SHH signaling.

## RESULTS

### Trisomy 21 diminishes Sonic Hedgehog signaling

To investigate how Trisomy 21 (T21) impacts SHH signaling, isogenic hTERT-RPE-1 cell lines with two (D21) or three (T21) copies of Hsa21 were studied (Lane et al., 2014). Ciliation was induced by serum depletion followed by SHH activation with the Smoothened agonist, SAG. SAG induction of SHH was initiated 24 hours after serum depletion and cells were analyzed 4 to 48 hours after SAG treatment which equates to 28 to 72 hours of serum depletion (Figure 1A). As previously observed, ciliation frequency was significantly reduced in T21 cells compared to D21 controls 24 hours after serum depletion (Figure 1B and Figure S1A; (Galati et al., 2018; McCurdy et al., 2022; Jewett et al., 2023)). By 36 hours of serum depletion, ciliation frequency of T21 cells matches that of D21 cells (Figure 1B, green).

**Figure 1:**
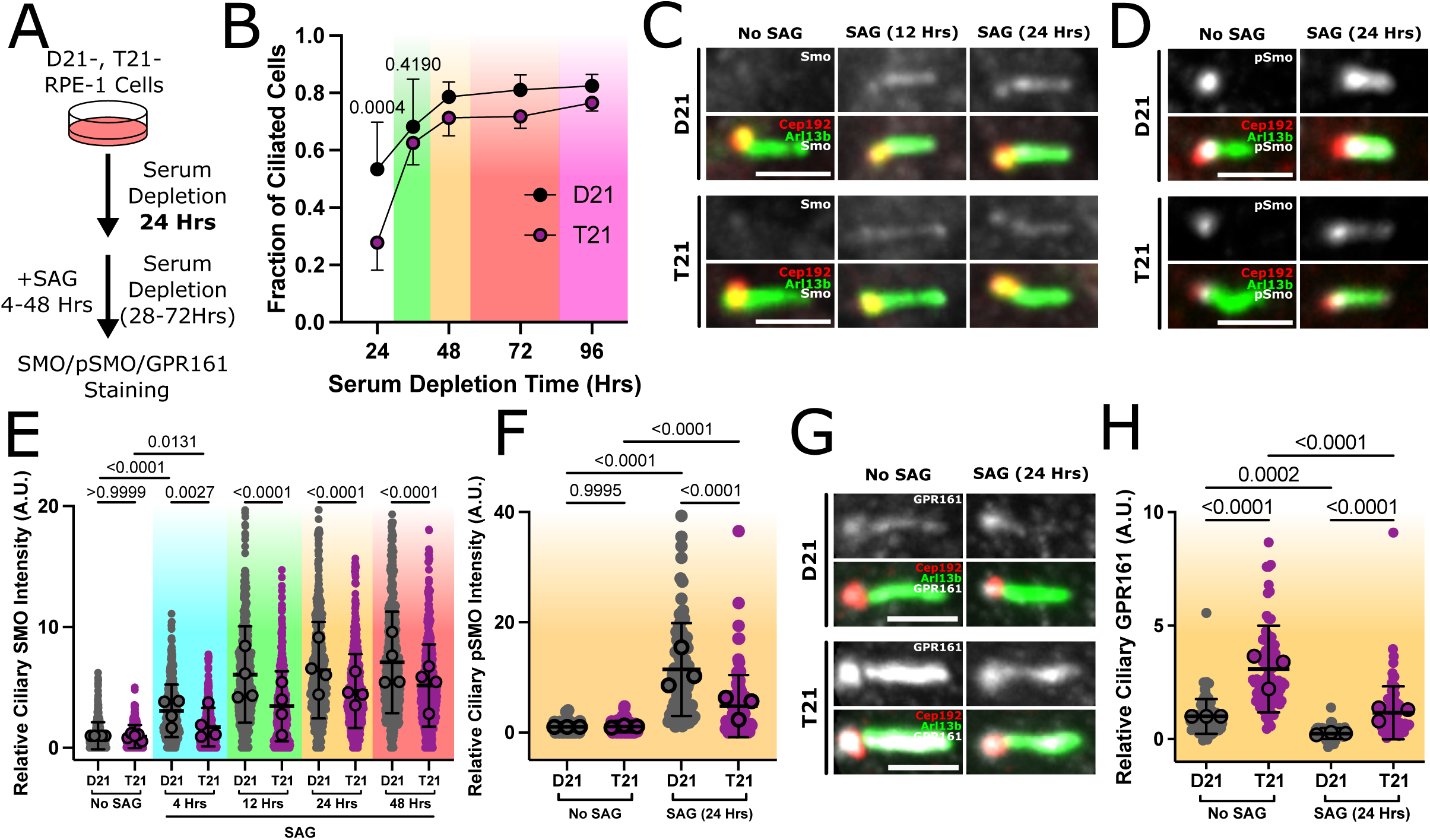
Trisomy 21 disrupts ciliary localization of proteins associated with SHH signaling. **(A)** Experimental design to determine whether Trisomy 21 affects ciliary localization of SMO. Isogenic disomy (D21) and trisomy 21 (T21) RPE-1 cell lines were cultured in serum-depleted medium for 24 hours to induce ciliation, treated with a Smoothened agonist (SAG) in serum-depleted medium for varying amounts of time (ranging from 28 – 72 hours total in serum depletion), and then fixed and stained for Smoothened (SMO), phosphorylated Smoothened (pSMO), and GPR161 protein. **(B)** T21 cells have fewer cilia compared to D21 cells 24 hours after serum depletion. Cilia frequencies in T21 approach those of D21 by 36 hours of serum depletion. Fraction of ciliated D21 and T21 cells along a time course of serum depletion. Color gradients indicate serum depletion time (24 hours, no color; 36 hours, green; 48 hours, yellow; 72 hours, red; 96 hours, magenta; n=4; 150-250 cells/replicate). **(C)** T21 decreases ciliary SMO levels compared to D21 cells. Representative confocal images of ciliary SMO in D21 and T21 cells at indicated times post SAG treatment. Cilia (ARL13b), green; centrosomes (CEP192), red; SMO, grayscale. Scale bar, 5µm. **(D)** T21 decreases ciliary pSMO levels compared to D21 cells. Representative confocal images of ciliary pSMO in D21 and T21 cells after 24hrs of SAG treatment (48 hours total serum depletion). Cilia (ARL13b), green; centrosomes (CEP192), red; pSMO, grayscale. Scale bar, 5µm. **(E)** Quantification of ciliary SMO intensity in D21 and T21 post SAG treatment. Color gradients indicate serum depletion time (24 hours, no color; 28 hours, cyan; 36 hours, green; 48 hours, yellow; 72 hours, red; n=4 biological replicates of 30-200 cilia/replicate). **(F)** Quantification of ciliary pSMO intensity with and without SAG treatment at 48 hours serum depletion (n=3; 30 cilia/replicate). **(G)** T21 increases ciliary GPR161 and diminishes SAG-induced GPR161 exit. Representative confocal images of ciliary GPR161 in D21 and T21 cells after 48 hours of serum depletion with or without SAG (24 hours). Cilia (ARL13b), green; centrosomes (CEP192), red; GPR161, grayscale. Scale bar, 5µm. **(H)** Quantification of ciliary GPR161 intensity in D21 and T21 cells after 48 hours of serum depletion with or without 24 hours of SAG treatment (n=3; 30 cilia/replicate). P-value, 2-Way ANOVA.

In ciliated cells, SMO levels in the primary cilium were measured to determine whether T21 affected SMO localization. SMO intensity in primary cilia was significantly reduced in T21 cells when compared to D21 after treatment with either SAG or SHH ligand (Figure 1C and 1E; Figure S1C and S1D). This reduction of SMO in T21 cilia persisted through the time course even after 36 hours of serum depletion when T21 ciliation frequency recovers to D21 levels (Figure 1B, green and Figure 1E, green). These differences were not due to variation in total cell SMO protein abundance between D21 and T21 cells (Figure S1B). Activation of SMO, which requires SMO phosphorylation (pSMO) mediates SHH signaling (Happ et al., 2022; Arveseth et al., 2021; Walker et al., 2024). In line with reduced ciliary SMO in T21 cells, pSMO levels were significantly decreased in T21 cilia compared to D21 cilia (Figure 1D and 1F; Figure S1F and S1G). Together, these observations suggest T21 cells exhibit diminished SMO localization and phosphorylation compared to D21 cells.

GPR161 is a constitutively active GPCR that localizes to the cilium in the absence of SHH activation (Mukhopadhyay et al., 2013). GPR161 suppresses SHH signaling by increasing cyclic AMP (cAMP) levels which promote Gli2/3 processing into its transcriptionally repressive form (GliR; (Mukhopadhyay et al., 2013)). Upon SHH activation, GPR161 decreases in the cilium in a SMO-dependent manner (Pal et al., 2016), thereby preventing Gli2/3 processing into GliR. After 48 hours of serum depletion, T21 cilia exhibit elevated levels of GPR161, regardless of SAG treatment, compared to D21 cells (Figure 1G and Figure 1H). These data suggest that T21 cells exhibited disrupted ciliary localization of multiple SHH-related proteins.

To test whether SHH transcriptional output is affected by T21, expression of SHH target genes, *GLI1* and *PTCH1*, was measured after SAG activation of SHH signaling. Both *GLI1* and *PTCH1* expression after SAG treatment were reduced in T21 cells compared to D21 (24 hours SAG and 48 hours total in serum depletion; Figure S1E). Interestingly, *PTCH1* expression is significantly increased in T21 cells without SAG treatment, compared to D21 cells. This is consistent with previous studies showing that neural progenitor cells from a murine Down syndrome model have increased *Ptch1* expression that suppresses SHH signaling and proliferation (Trazzi et al., 2011). Although SAG directly activates SMO independent of PTCH (Chen et al., 2002), it is possible the reduction of ciliary SMO in these T21 cells is due, in part, to increased *PTCH1* expression and ciliary localization. However, the reduction of ciliary SMO in T21 cells compared to D21 after treatment with purified SHH ligand (SHH-N) suggests SMO localization to T21 cilia may be independent of PTCH, if PTCH ciliary exit is similar between T21 and D21 cells (Figure S1C and S1D). Unfortunately, we were unable to visualize endogenous PTCH protein to determine whether T21 cells display altered ciliary localization or exit of PTCH. Collectively, these data suggest that even though T21 cells eventually construct primary cilia, they do not effectively signal SHH.

### Diminished ciliary Smoothened in T21 is not due to increased Pericentrin

Delayed primary ciliogenesis in T21 cells results from elevated PCNT which is proposed to impede the transport of cilia assembly factors to the centrosome (Galati et al., 2018; McCurdy et al., 2022; Jewett et al., 2023). Localization of ciliary proteins to the centrosome and subsequent ciliogenesis are rescued in T21 when PCNT is reduced to disomic levels. To determine whether reduced PCNT levels could also rescue ciliary SMO levels in T21 cells, PCNT siRNA was used to knock down PCNT to nearly disomic and sub-disomic levels (Figure 2A-C and Figure S2A). T21 ciliation frequency is partially rescued when PCNT is reduced below D21 levels (Figure 2D and Figure S2B; (McCurdy et al., 2022)). Ciliary SMO levels in T21 cells remain significantly decreased compared to D21 cells when PCNT is reduced to near or below D21 levels (Figure 2E and 2F). Moreover, T21 cells with two copies of *PCNT* after CRISPR-Cas9 deletion of one copy (T21-T3D2 cells) also exhibit significantly reduced levels of ciliary SMO compared to D21 cells (Figure S2E and S2F). These cells exhibit sub-disomic levels of PCNT protein (Figure S2C and S2D). Finally, when comparing ciliary SMO to centrosomal/pericentrosomal PCNT on a cell-by-cell basis, increased PCNT was not correlated with decreased ciliary SMO (Figure S2G). Overall, data from both siRNA-mediated depletion experiments and T21-T3D2 cells suggest impeded SMO localization to T21 primary cilia is not due to increased PCNT. This is in contrast to wild-type human primary fibroblast cultures where overexpression of PCNT to T21 levels was sufficient to reduce ciliary SMO (Galati et al., 2018). Thus, SMO ciliary localization likely depends on multiple factors; while acute PCNT overexpression is sufficient to reduce ciliary SMO, possibly because cells lack the compensatory mechanisms to accommodate a sudden increase, chronically reducing PCNT to near- and sub-disomic levels is insufficient to rescue the decrease in ciliary localization of SMO in T21. This suggests that additional T21-associated centrosomal/pericentrosomal factors could contribute to SMO mislocalization.

**Figure 2:**
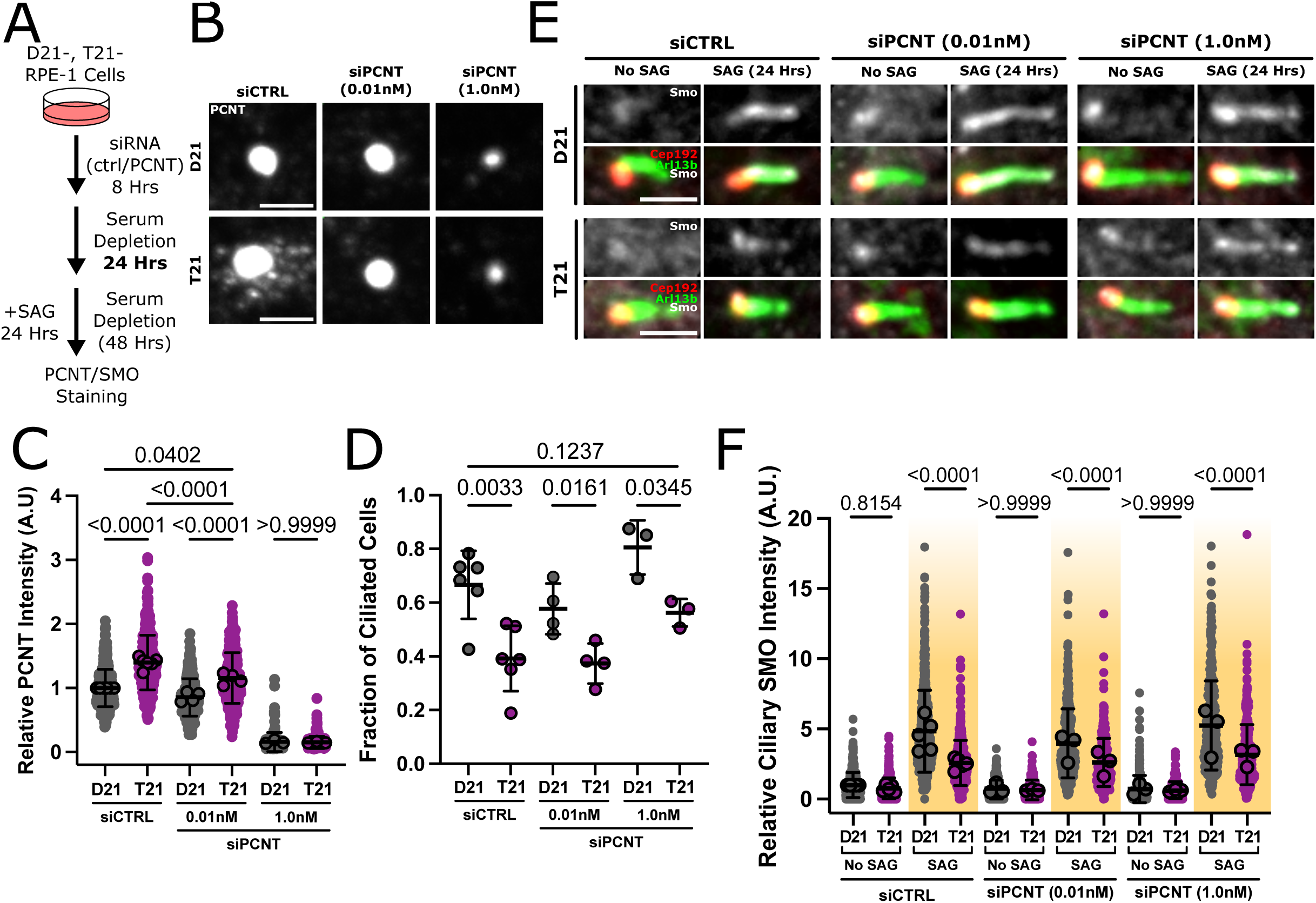
Elevated PCNT in Trisomy 21 does not affect ciliary SMO. **(A)** Experimental design to determine whether increased Pericentrin (PCNT) in T21 cells affects SMO localization to the cilium. Isogenic RPE-1 D21 and T21 cells were treated with control or PCNT siRNA for 8 hours, serum depleted for 24 hours to induce ciliation, and treated with SAG for 24 hours in serum-depleted medium (48 hours total in serum depletion). Cells were then fixed and stained for PCNT or SMO. **(B)** Representative confocal images of PCNT knock down to approximately disomic (0.01nM siPCNT) and sub-disomic (1.0 nM siPCNT) levels in D21 and T21 cells. PCNT, grayscale. Scale bar, 5µm. **(C)** Quantification of PCNT levels at the centrosome in control and siPCNT-treated D21 and T21 cells after 24 hours of serum depletion. **(D)** Ciliation increases with depletion of PCNT to sub disomic levels. Quantification of ciliation frequency in control and siPCNT-treated D21 and T21 cells after 24 hours of serum depletion. **(E)** Representative confocal images of ciliary SMO in control and siPCNT-treated D21 and T21 cells after 24 hours of SAG treatment (48 hours total in serum depletion). Cilia (ARL13b), green; centrosomes (CEP192), red; SMO, grayscale. Scale bar, 5µm. **(F)** Reduction of PCNT does not rescue ciliary SMO localization in T21 cells. Quantification of ciliary SMO intensity in control and siPCNT-treated D21 and T21 cells with and without SAG treatment. Color gradients indicate serum depletion times (24 hours, no color; 48 hours, yellow; n=4; 30-130 cilia/replicate). P-value, 2-Way ANOVA.

### Smoothened protein localization to centrosomes is unaffected in Trisomy 21

Before proteins enter the cilium, they first transport to the centrosome (Wheway et al., 2018). Given that some ciliary proteins mislocalize around the centrosome in T21 (McCurdy et al., 2022; Galati et al., 2018; Jewett et al., 2023), we examined whether T21 alters SMO protein localization at the centrosome (Figure 3A and 3B). A moderate increase in SMO protein was observed at T21 centrosomes compared to D21 centrosomes before SAG treatment (24 hours of serum depletion). However, no significant difference in centrosome-localized SMO intensity was observed between D21 and T21 cells, regardless of the duration of SAG treatment (4-48 hours of SAG treatment and 28–72 hours total in serum depletion; Figure 3A and 3B).

**Figure 3:**
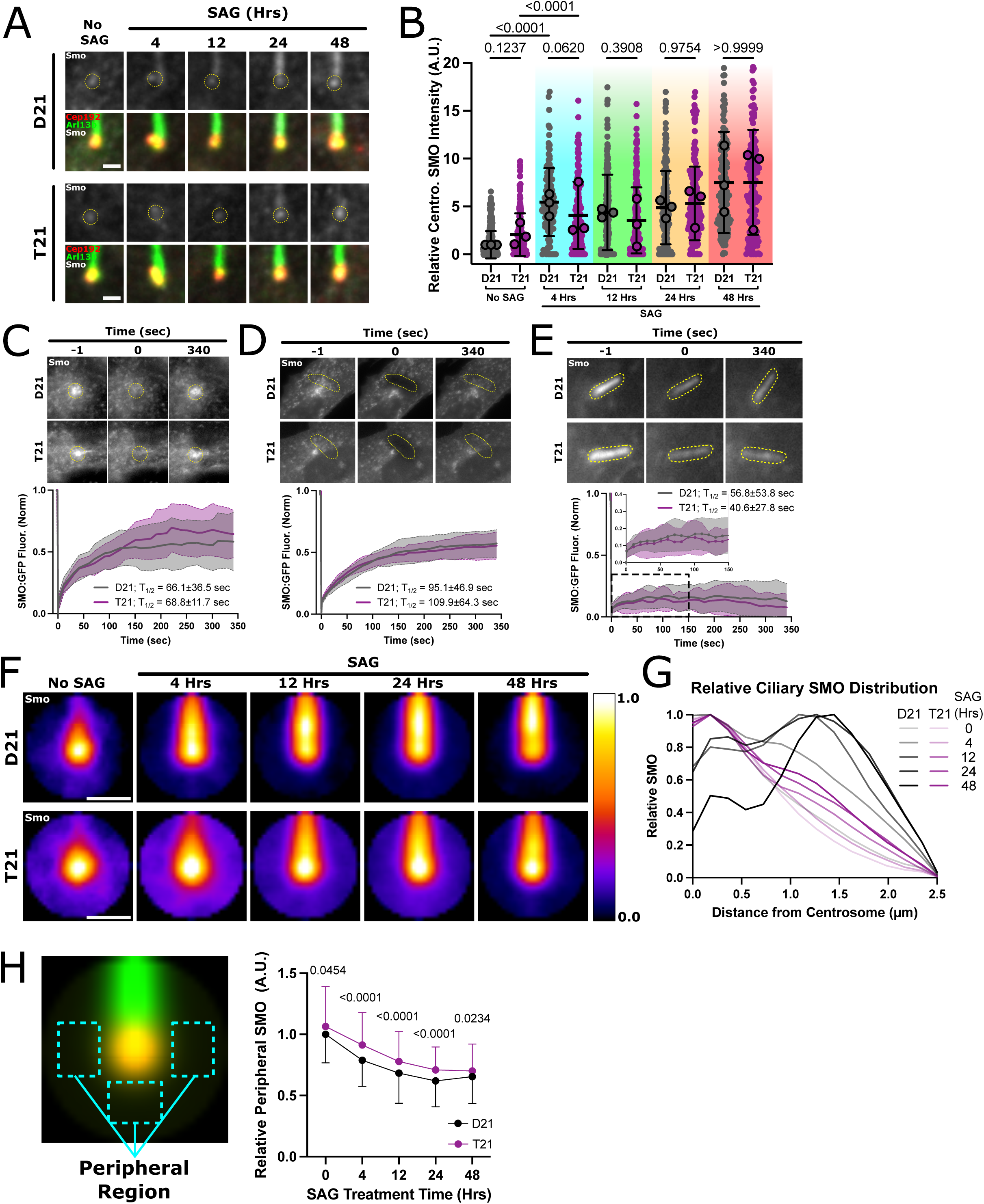
SMO localization to the centrosome is unchanged in Trisomy 21. **(A)** Representative confocal images of SMO localization at the centrosome in D21 and T21 cells across a SAG treatment time course. Cilia (ARL13b), green; centrosomes (CEP192), red; SMO, grayscale. Circles indicate regions that were quantified. Scale bar, 2µm. **(B)** Quantification of centrosome-localized SMO intensity in a 2.5µm diameter region centered on CEP192-labeled centrioles. Color gradients indicate total time in serum depletion (24 hours, no color; 28 hours, cyan; 36 hours, green; 48 hours, yellow; 72 hours, red; n=3; 30-90 centrosomes/replicate; P-value, 2-Way ANOVA). **(C-E)** SMO turnover is unchanged in T21 at the centrosome and peripheral regions, and cilia have a stable pool of SMO protein. Representative FRAP images and corresponding quantification showing SMO turnover in SMO:GFP expressing D21 and T21 cells at the centrosome (C), peripheral regions (D), and cilium (E). T_1/2_ values were calculated using EasyFRAP webtool. P-Value, *t*-test. **(F)** T21 reduces ciliary SMO and increases accumulation of centrosomal SMO. Image averages of cropped and oriented cilia stained for SMO. White/Yellow colors indicate elevated SMO localization while Black/Blue colors indicate lower SMO localization. (n=3; 100-300 images/replicate). Scale bar, 2µm. **(G)** T21 restricts SMO to the ciliary base. Line scan analysis of SMO distribution along cilia (n=3; 30 cilia/replicate). **(H)** SMO protein persists longer in peripheral regions in T21 cells compared to D21 cells. SMO protein localization was measured and plotted after image averaging. Example of peripheral regions is depicted.

We next tested whether the rate of SMO turnover at centrosomes was altered in T21. Fluorescence recovery after photobleaching (FRAP) was used to measure stably overexpressed SMO:GFP protein turnover after 4 hours of SAG treatment in D21 and T21 cells. The SMO:GFP recovery rate at T21 centrosomes was similar to D21 (Figure 3C; T_1/2_ = D21: 66.1±36.5 seconds; T21: 68.8±11.7 seconds; *p*=0.9106). Total SMO:GFP recovery at T21 centrosomes was elevated but not significantly different from D21 (Figure 3C; percent recovery = D21: 58.9±22.7%; T21: 69.6±19.5%; *p*=0.1283). This may reflect variability in the exchange of SMO protein between centrosome peripheral pools and the centrosome. These results suggest that both the rate and extent of SMO:GFP turnover at the centrosome are variable yet comparable between D21 and T21 cells.

To determine whether SMO turnover dynamics surrounding the centrosome were altered in T21, overexpressed SMO:GFP was photobleached in regions peripheral to the centrosome and fluorescence recovery was measured (Figure 3D). These regions were selected so that photobleaching occurred as close as possible to the centrosome without including the centrosome itself. SMO:GFP recovery dynamics and total recovery at these regions were similar between D21 and T21 cells (Figure 3D; T_1/2_ = D21: 95.1±46.9 seconds; T21: 109.9±64.3 seconds; *p*=0.4340; percent recovery = D21: 57.3±11.1%; T21: 55.5±11.0%; *p*=0.6810). These data suggest that overexpressed SMO:GFP turnover at and around the centrosome is comparable between D21 and T21 cells.

To test whether SMO protein dynamics within the cilium were altered in T21 cells, SMO:GFP turnover dynamics were measured using FRAP in D21 and T21 cilia. Consistent with prior studies, low SMO:GFP percent recovery was observed in primary cilia of both D21 and T21 cells (Figure 3E; (Dyson et al., 2016)). Moreover, no significant differences were observed in SMO:GFP recovery between D21 and T21 cilia (Figure 3E; T_1/2_ = D21: 56.8±53.8 seconds; T21: 40.6±27.8 seconds; p=0.6756; percent recovery = D21: 16.9±12.3%; T21: 14.8±9.6%; *p*=0.6054). Thus, no measurable difference in low SMO:GFP recovery was observed between D21 and T21 cilia.

To capture the low-level changes in SMO protein localization at and around the cilium, a time course of SMO protein localization was visualized in image averages after SAG treatment (Figure 3F). In line with previous results (Figure 1E), SMO localized to the proximal end of the T21 cilium and was more uniformly distributed in D21 cilia (Figure 3F and 3G). This suggests SMO movement throughout the cilium was disrupted in T21 compared to D21. In addition, SMO protein persisted peripheral to the centrosome after SAG treatment in T21 cells as noted by elevated peripheral SMO at 4 hours (28 hours total of serum depletion) in T21 but not in D21 (Figure 3F and 3H). These data suggest SMO protein enters and moves within primary cilia less efficiently within T21 cells compared to D21. The persistence of SMO protein peripheral to the centrosome in T21 cells suggests T21 may limit SMO protein transport from the centrosome and pericentrosomal region(s) to the cilium (Figure 3F-3H).

In summary, SMO localization and turnover at centrosomes appears to be unchanged in T21 cells compared to D21. The reduced levels of ciliary SMO may be attributed to defects in entry or transport within the cilium that were undetected in our SMO protein dynamics studies.

### Cilia maturation equilibrates ciliary Smoothened levels and GPR161 ciliary exit between D21 and T21

In T21 cells, primary ciliogenesis is delayed such that, given enough time to ciliate, T21 cells form a primary cilium at the same frequency as D21 cells (Figure 1B; (Galati et al., 2018; McCurdy et al., 2022; Jewett et al., 2023)). To determine whether mature T21 primary cilia become competent to localize SMO at levels like D21, the duration of serum depletion and therefore cilia maturation was increased to 72 hours followed by 24 hours of SAG treatment (96 hours total serum depletion; Figure 4A). With extended ciliary maturation, SAG-induced SMO accumulation in T21 cilia is comparable to D21 (Figure 4B and 4D, magenta). Normalization of ciliary SMO levels between D21 and T21 cells was not due to changes in PCNT levels (Figure S3A and S3B). Importantly, ciliary SMO levels in D21 were also decreased in these experiments compared to 48 hours of total serum depletion (Figure 1E, yellow and Figure 4D, magenta). Thus, the equilibration results from decreased SMO levels in mature D21 cilia. Regardless, this suggests that when given enough time to mature, primary cilia recruit comparable SMO levels between D21 and T21 upon SAG treatment.

**Figure 4:**
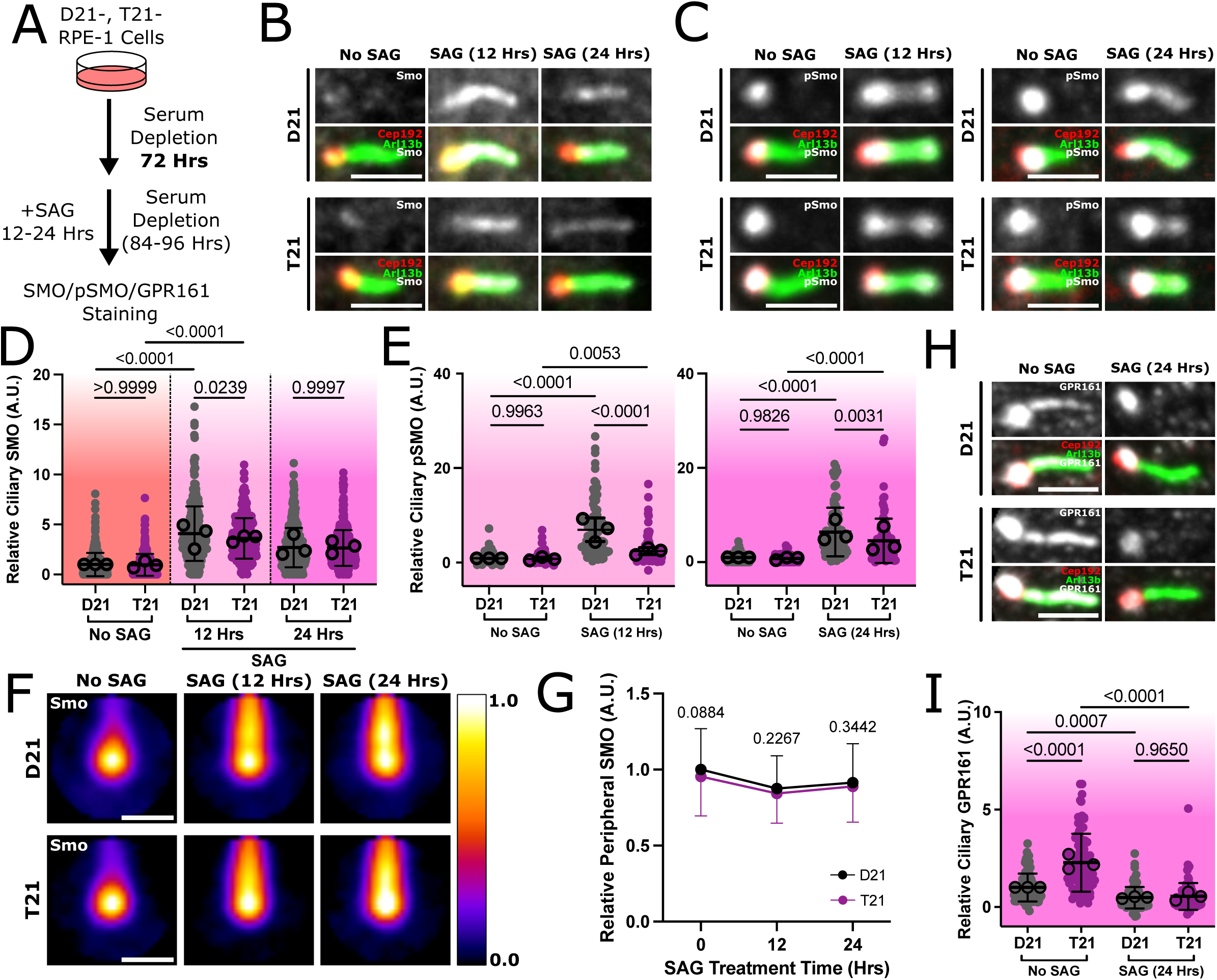
Cilia maturation equalizes ciliary localization of SHH-associated proteins between D21 and T21. **(A)** Experimental design to determine whether reduced ciliary SMO localization in T21 cells can increase with longer cilia maturation time. Isogenic RPE-1 D21 and T21 cells were cultured in serum depleted medium for 72 hours to induce ciliation and cilia maturation. Cells were then treated with SAG in serum-depleted medium before fixation and staining for SMO and pSMO (72 – 96 hours total in serum depletion). **(B)** Ciliary SMO levels are similar between T21 and D21 cells with increased cilia maturation time. Representative confocal images of ciliary SMO post serum depletion and indicated SAG treatment duration. Cilia (ARL13b), green; centrosomes (CEP192), red; SMO, grayscale. Scale bar, 5µm. **(C)** Ciliary pSMO levels are partially rescued between T21 and D21 cells with increased cilia maturation time. Representative confocal images of ciliary pSMO at 96 hours of serum depletion with or without SAG (24 hours). Cilia (ARL13b), green; centrosomes (CEP192), red; pSMO, grayscale. Scale bar, 5µm. **(D)** Quantification of ciliary SMO intensity in D21 and T21 cells at indicated times post SAG treatment. Color gradients indicate serum depletion time (72 hours, red; 84 hours, salmon; 96 hours, magenta; n=3; 90-150 cilia/replicate). **(E)** Quantification of ciliary pSMO intensity in D21 and T21 cells at 84 hours of serum depletion with or without 12 hours of SAG treatment and 96 hours of serum depletion with or without 24 hours of SAG treatment (n=3; 30 cilia/replicate). **(F)** Increased ciliary maturation time rescues differences in ciliary and centrosomal SMO localization. Image averages of cropped and oriented cilia stained for SMO. White/Yellow colors indicate elevated SMO localization while Black/Blue colors indicate lower SMO localization. (n=3; 90-150 cilia/replicate; averages of >300 cilia). **(G)** Peripheral SMO localization was measured and plotted after image averaging. Example of peripheral regions is depicted in Figure 3H. **(H)** SAG-induced GPR161 exit is rescued by cilia maturation. Representative confocal images of ciliary GPR161 in D21 and T21 cells after 96 hours of serum depletion with or without SAG (24 hours). Cilia (ARL13b), green; centrosomes (CEP192), red; GPR161, grayscale. Scale bar, 5µm. **(I)** Quantification of ciliary GPR161 intensity in D21 and T21 cells with or without SAG treatment. Color gradients indicate serum depletion times (96 hours, magenta; n=3; 30 cilia/replicate). P-value, 2-Way ANOVA.

To determine whether phosphorylated SMO in T21 cells was also increased with prolonged cilia maturation, ciliary pSMO levels were measured. Despite similar total SMO levels in D21 and T21 cilia, ciliary pSMO levels remained significantly reduced in T21 cells compared to D21 (Figure 4C and 4E). However, the ratio of pSMO intensity between T21 and D21 cilia at mature cilia at 96 hours of serum depletion (T21/D21 = 0.67±0.15; Figure 4E) is substantially greater than the ratio at early cilia at 48 hours of total serum depletion (T21/D21 = 0.41±0.14; *p*=0.0968; Figure 1F). In summary, increased cilia maturation can equilibrate SMO ciliary localization and partially equilibrate SMO phosphorylation in D21 and T21 cilia.

To capture the slow changes in SMO protein localization at and around the cilium, a time course of SMO protein localization was visualized in image averages after SAG treatment with increased cilia maturation. Similar to 48 hours of serum depletion, SMO localized to the proximal end of the cilium in T21 while D21 SMO was more uniformly distributed throughout the cilium (Figure S3C). This suggests that despite having similar amounts of SMO in mature D21 and T21 cilia, SMO localization remains aberrant in T21 cells compared to D21 cells. SMO localization around the centrosome was similar between D21 and T21 cells in mature cilia (Figure 4F and 4G), which contrasts less mature cilia (Figure 3F and 3H). These data suggest that increased cilia maturation may improve SMO entry into the cilium but that movement within the cilium remains disrupted in T21 cilia compared to D21.

Because the abnormal localization of SMO in T21 cilia can be equalized to D21 by allowing more time for cilia maturation, the localization of other SHH-associated proteins was also examined after extending cilia maturation time. The IFT-B complex protein, IFT88, promotes anterograde transport of cargo along the cilia axoneme. IFT88 is postulated to facilitate SMO transport along the cilium (Haycraft et al., 2007; Wang et al., 2009). IFT88 levels were assessed in D21 and T21 cilia to determine if diminished ciliary SMO could be explained by changes to IFT88 localization. Surprisingly, IFT88 levels are increased in T21 cilia compared to D21 after 24 hours of serum depletion (Figure S3D and Figure S3E, no color). IFT88 levels in the primary cilium increase with maturation, equilibrating between D21 and T21 by 48 hours of serum depletion (Figure S3D and S3E, yellow). By 72 hours of serum depletion, IFT88 levels in T21 are modestly, but significantly, decreased compared to D21 (Figure S3D and S3E, red). To determine whether equilibrated IFT88 levels coincide with relative changes to SMO localization along cilia, IFT88 and SMO localization along D21 and T21 cilia were compared at 48 hours of serum depletion with 24 hours of SAG treatment. Despite similar enrichment of IFT88 at the ciliary base in both D21 and T21 cells, SMO localization remains focused to the proximal end of T21 cilia (Figure S3F and Figure 3G). These data suggest that altered ciliary SMO distribution in T21 cells is not due to changes in IFT88 localization. However, IFT trains that interact with SMO may have disrupted movement along the primary cilium in T21 cells. Indeed, disrupted IFT movement along T21 cilia could explain the proximal localization of ciliary SMO in T21 cells and may not be affected by ciliary maturation (Figure 3G; Figure S3C).

Finally, to determine whether increased cilia maturation could normalize GPR161 localization, ciliary GPR161 levels were measured after increasing serum depletion time. After 96 hours of serum depletion, GPR161 remained elevated in T21 cilia, compared to D21 cilia, without SAG treatment (Figure 4H and 4I). However, levels of GPR161 are comparable between D21 and T21 after SAG treatment in mature cilia (Figure 4H and 4I), suggesting that increased cilia maturation can rescue ciliary GPR161 exit in T21 cells. In summary these data suggest that while ciliary localization of SHH signaling proteins is altered in early or immature T21 cilia, localization of these proteins is similar to D21 in mature cilia. While delayed maturation may alter ciliary localization of SHH proteins in T21 cells, whether activity of these proteins is altered and subsequently disrupts SHH signaling remains to be determined. Thus, these data suggest that delayed maturation of primary cilia in T21 may contribute to defective SHH signaling.

## DISCUSSION

This study expands upon our prior discovery that T21 delays primary ciliogenesis and reduces downstream SHH responses (Galati et al., 2018; Jewett et al., 2023; Klein et al., 2021; Roper et al., 2006; Moyer et al., 2023). Decreased SMO localization to the cilium in T21 does not appear to arise from elevated PCNT or defective recruitment to centrosomes, but instead, defective localization and activation of SMO in the cilium. Interestingly, if cilia are provided with more time for maturation, ciliary SMO levels equilibrate between D21 and T21 cells. Thus, T21 disruption of SHH signaling may arise, in part, through delayed cilia maturation.

Our findings echo previous reports that T21 delays primary ciliogenesis and reduces levels of ciliary SMO protein (Figure 1; (Galati et al., 2018; Jewett et al., 2023)) and provide mechanistic insights into how T21 may diminish SHH signaling. First, SMO phosphorylation (pSMO), which is critical for SMO activation, is impaired in T21 cilia (Figure 1F). SMO activation promotes SHH signaling, in part, by sequestering PKA from Gli2/3 (Happ et al., 2022; Arveseth et al., 2021; Walker et al., 2024). Second, the SHH suppressor GPR161 protein is elevated in T21 primary cilia. Normally, activation of SHH signaling by SAG causes depletion of GPR161 from the cilium (Pal et al., 2016). However, this is disrupted in T21 and GPR161 levels remain elevated (Figure 1G and 1H). GPR161 suppresses SHH signaling by constitutively producing cAMP, thus promoting PKA-mediated processing of Gli2/3 transcription factors into repressive forms (GliR; (Mukhopadhyay et al., 2013)). Thus, one possible model is that T21 suppresses SHH signaling through synergistic events, such as increased ciliary GPR161 and diminished SMO activation, that lead to increased levels of available cAMP and subsequently promote Gli2/3 conversion to GliR.

SMO activation and GPR161 ciliary exit are both dependent on SMO retention in the cilium (Walker et al., 2024; Pal et al., 2016). Because we observe reduced ciliary localization of SMO in T21 without changes to centrosome recruitment, this suggests that diminished SMO localization to primary cilia in T21 is due to a defect in SMO entry or retention in cilia, rather than a defect in SMO transport to centrosomes (Figure 3). Alternatively, SMO ciliary mislocalization in T21 cells could be due to alterations in SMO ubiquitination dynamics that affect localization or destruction of SMO protein (Desai et al., 2020; Shinde et al., 2020; Lv et al., 2021, 2022). Indeed, the altered SMO localization observed in T21 cells (Figure 3F-3H) could be due to increased SMO ubiquitination that impedes proper localization to the cilium when stimulated with SAG. In either model, reduction of ciliary SMO may lead to the perturbations in pSMO and GPR161 levels we observe in T21 cilia (Walker et al., 2024; Pal et al., 2016). However, the ciliary defects that suppress ciliary SMO accumulation remain to be understood.

Ciliary PTCH can also recruit E3 ubiquitin ligases to the cilium to alter SMO localization and trigger SMO proteolysis (Lv et al., 2021; Desai et al., 2020; Shinde et al., 2020). A role for ciliary PTCH protein in mediating the reduced ciliary SMO levels in T21 cannot be ruled out. Our data and others suggest that *PTCH1* expression is upregulated in T21, resulting in more ciliary PTCH protein (Figure S1E; (Trazzi et al., 2011)). However, SAG directly interacts with SMO independent of PTCH, and treating cells with purified SHH ligand (SHH-N) similarly resulted in reduced ciliary SMO in T21 cells (Figure S1C and S1D). This suggests that the reduced ciliary SMO levels in T21 are unlikely to be explained solely by elevated ciliary PTCH protein levels. However, we were unable to detect whether more PTCH protein localizes to the primary cilium in T21 cells and impedes SMO accumulation in the cilium.

Prolonged serum depletion is sufficient for T21 cells to form mature primary cilia with SMO levels and partially rescued SMO activation comparable to D21 cells (Figure 4). Moreover, increased cilia maturation equalizes SMO localization at regions peripheral to the centrosome (Figure 4F and 4G) and is associated with SAG-induced reduction in GPR161 (Figure 4H and 4I). One potential model is that cilia maturation normalizes SMO ubiquitination dynamics in T21 cells, thereby correcting SMO and GPR161 ciliary localization as described above (Desai et al., 2020; Shinde et al., 2020; Lv et al., 2021, 2022; Pal et al., 2016). An alternative model is that T21 cells exhibit altered ciliary membrane phospholipid composition that is subsequently corrected with cilia maturation. SMO and GPR161 localization to cilia are modulated by ciliary phosphoinositide content (Dyson et al., 2016; Chávez et al., 2015; Garcia-Gonzalo et al., 2015). Elevated phosphatidylinositol 4,5-bisphosphate (PI(4,5)P_2_) promotes GPR161 localization to the cilium, while elevated ciliary phosphatidylinositol 4-phosphate (PI(4)P) may promote SMO retention in primary cilia, though results vary across reports (Dyson et al., 2016; Chávez et al., 2015; Garcia-Gonzalo et al., 2015; Jiang et al., 2016; Kumari et al., 2022). Thus, ciliary maturation may involve corrections to the ciliary phospholipid composition in T21 cells that subsequently correct SMO and GPR161 localization. More studies are needed to further define features of cilia maturation and determine how T21 may impede them.

Delayed primary cilia formation and SHH signaling in T21 likely disrupts SHH responses during development (Galati et al., 2018; Jewett et al., 2023). Under most developmental contexts, SHH signal transduction depends on duration and amount of SHH ligand available, which are under tight temporal control to guide proliferation and differentiation dynamics during development of tissues and organs, such as the brain and heart (Groves et al., 2020; Zhang and Beachy, 2023; Nguyen et al., 2022). For example, cerebellar granule cells and cardiac progenitors must proliferate and differentiate at appropriate times to reach sufficient cerebellar volume and complexity in the brain and to form specialized cardiac structures in the heart (Roper et al., 2006; Jewett et al., 2023; Klein et al., 2021; Smeyne et al., 1995; Wechsler-Reya and Scott, 1999; Rowton et al., 2022). Although cilia maturation normalizes ciliary localization of SHH proteins in T21, the timing of SHH ligand exposure in development may not permit more time for these cells to mature their cilia and respond effectively. Therefore, T21-dependent delays in cilia formation and maturation could manifest in the tissue-level phenotypic changes typically observed in Down syndrome.

### Limitations of the study

There are several limitations of this study that should also be considered when evaluating and interpreting the data presented here. First, the study is limited to the use of isogenic RPE-1 cells lines. As a result, the altered localization of SHH proteins shown in T21 cells may not be representative of other T21 models, such as murine models of Down syndrome or primary patient samples. Moreover, although these data demonstrate that T21 alters the localization of several proteins involved in SHH signaling, the functional activity of these proteins in T21 cells was not assessed. Decreased localization of specific proteins to T21 cilia may be compensated by increased functional activity, and vice versa. Furthermore, the protein dynamics studies rely on exogenous overexpression of SMO:GFP. Exogenously overexpressed SMO is known to accumulate in primary cilia regardless of SHH stimulation (Kovacs et al., 2008). Therefore, the similar SMO turnover rates between D21 and T21 cells may reflect the tendency of the overexpressed SMO protein to accumulate in primary cilia, independent of potential T21-associated defects.

## MATERIALS AND METHODS

### Cell Culture

Disomy 21 (D21) and trisomy 21 (T21) hTERT-immortalized retinal pigment epithelial (RPE-1) cells were generated by Andrew Lane and David Pellman (Lane et al., 2014). Cells were grown in DMEM/F12 (11330-032; Gibco) supplemented with 10% fetal bovine serum (FBS; Phoenix Scientific; PS-100) and 1% penicillin/streptomycin at 37°C and 5% CO_2_. Cells were passaged at 1:10 at 80-90% confluency using 0.25% trypsin (15090-046; Gibco). Cells were screened for mycoplasma every 6 months.

### Immunofluorescence

12mm glass coverslips were washed in 1M HCl heated to 50°C for 16 hours. Coverslips were then washed with water, followed by 50%, 70%, and 95% ethanol in a sonicating water bath for 30 minutes. Coverslips were coated with collagen (C9791; Sigma) and left to air dry for 15 minutes prior to crosslinking under UV light for an additional 15 minutes. Cover slips were then washed three times with phosphate buffered saline (PBS; 1 mM KH_2_PO_4_, 155 mM NaCl, 3 mM Na_2_-HPO_4_-7H_2_O, pH 7.4) before plating cells. For siRNA experiments, 40,000 cells were plated on each coverslip. For all other experiments, 50,000 – 60,000 cells were plated on each coverslip. To induce ciliation, and as a model for cilia maturation, cells were cultured in serum-depleted medium (DMEM/F12 (11330-032; Gibco) supplemented with 0.5% fetal bovine serum (FBS; Phoenix Scientific; PS-100) and 1% penicillin/streptomycin) for a minimum of 24 hours prior to treatment with 200nM Smoothened agonist (SAG; SML1314; Sigma). For experiments in Figure 1E, Figure 3A-3B, and Figure 3F-3H, cells were treated with SAG at 24 hours of serum depletion, and the total time in serum depletion ranged from 24 hours (no SAG) to 72 hours (48 hours of SAG). For experiments in Figure 1C-1H and Figure 2E and 2F, cells were treated with SAG at 24 hours of serum depletion, and the total time in serum depletion was 48 hours (24 hours in SAG). For experiments in Figure 4, cells were cultured in serum-depleted (0.5% FBS) medium for 72 hours to facilitate cilia maturation prior to SAG treatment, and the total time in serum depletion ranged from 72 hours (no SAG) to 96 hours (24 hours of SAG). For experiments in Figure S3A-S3B and S3D-S3E, cells were cultured in serum-depleted (0.5% FBS) medium for 24 hours, 48 hours, or 72 hours prior to fixation. For experiments using GPR161 and phospho-SMO antibodies, cells were fixed using 4% paraformaldehyde in PBS (15710; Electron Microscopy Sciences) for 15 minutes at room temperature followed by fixation/permeabilization in ice cold 100% methanol for 3 minutes at -20°C. Coverslips were washed 3 times for 5 minutes each in PBS then stored in PBS at 4°C until immunostaining. For experiments using SMO and IFT88 antibodies, cells were fixed with ice cold methanol for 10 minutes at -20°C. Coverslips were washed three times for 5 minutes each in PBS and stored in PBS at 4°C until permeabilization and subsequent immunostaining. Fixed cells were permeabilized with 0.5% Triton X-100 in PBS for 10 minutes at room temperature then washed three times for 5 minutes each in PBS. For immunostaining in all experiments, fixed cells were blocked with 0.5% bovine serum albumin (BSA; A3912; Sigma) in PBS with 0.5% NP-40 (IGEPAL, 18896; Sigma), 1 mM MgCl_2_ and 1 mM NaN_3_ for 1 hour at room temperature. Cells were incubated overnight in primary antibodies diluted in blocking buffer: mouse anti-Smoothened (1:500; sc-166685; Santa Cruz Biotechnologies), rabbit anti-ARL13B (1:500; 177-1-AP; ProteinTech), mouse anti-ARL13B (1:300; N295B/66; Developmental Studies Hybridoma Bank, University of Iowa), goat anti-CEP192 (1:2000; gift from Andrew Holland’s lab), rabbit anti-Phospho-Smoothened (1:300; 7TM0239A-IC; 7TM Antibodies), rabbit anti-GPR161 (1:300; 29328-1-AP; ProteinTech), rabbit anti-IFT88 (1:200; 13967-1-AP; ProteinTech), and rabbit anti-Pericentrin (1:2000; ab4448; Abcam). Coverslips were washed three times for 5 minutes each in PBS then incubated for 1 hour at room temperature in secondary antibodies and Hoechst-33258 (1µg/mL; Invitrogen) diluted in blocking buffer: AlexaFluor-488 donkey anti-rabbit (1:1000; A21206; Invitrogen), AlexaFlour-647 donkey anti-mouse (1:1000; A31571; Invitrogen), and AlexaFluor-594 donkey anti-goat (1:1000; A11058; Invitrogen). Coverslips were washed four times for 5 minutes each in PBS before mounting to slides using Citifluor (17970-25; Electron Microscopy Sciences). Coverslips were sealed to slides using clear nail polish.

### RT-PCR

D21- and T21-RPE-1 cells were plated on 6cm dishes (340,000 per dish) in DMEM/F12 (11330-032; Gibco) supplemented with 10% fetal bovine serum (FBS; Phoenix Scientific; PS-100) and 1% penicillin/streptomycin and incubated for 24 hours at 37°C and 5% CO_2_. Cells were cultured in DMEM/F12 (11330-032; Gibco) supplemented with 0.5% fetal bovine serum (FBS; Phoenix Scientific; PS-100) and 1% penicillin/streptomycin for 48 hours. After 24 hours of serum depletion, cells were treated with 1µM SAG (SML1314; Sigma) or vehicle control for 24 hours. Cells were collected from dishes using 0.25% trypsin (15090-046; Gibco) and pelleted prior to RNA isolation. RNA was isolated using Quick^TM^ DNA/RNA MiniPrep Kit (ZD7001; Zymo) according to manufacturer’s protocol. cDNA was synthesized from isolated RNA using SuperScript^TM^ IV First-Strand Synthesis System (18091050; Invitrogen) according to manufacturer’s protocol. All PCR reactions were performed using Phusion Polymerase and Phusion HF Buffer (B0518S; New England Biolabs). Oligonucleotides used in PCR reactions consist of Gli1 (F: 5’ – CCCAGTACATGCTGGTGGTT – 3’; R: 5’ – GCTTTACTGCAGCCCTCGT – 3’), Ptch1 (F: 5’ – GGCAGCGGTAGTAGTGGTGT – 3’; R: 5’ – TGTAGCGGGTATTGTCGTGTG – 3’), and ß-Actin (F: 5’ - GAAAATCTGGCACCACACCT – 3’; R: 5’ – TAGCACAGCCTGGATAGCAA – 3’). PCR products were run on 1.2% Agarose gel, stained with Ethidium Bromide, and visualized using Gel Doc^TM^ EZ Imager (170-8270; BioRad). Bands were quantified using ImageJ.

### RNAi

Human PCNT siRNA (Smart Pool; M-012172-01-0005; Dharmacon) was transfected into cells using lipofectamine RNAi MAX (13778100; ThermoFisher Scientific) according to the manufacturer’s protocol. Final concentration for Human PCNT siRNA ranged from 0.01 – 1 nM (as indicated in Figure 2). Mission siRNA universal negative control #1 (SIC001-1NMOL; Sigma) was used at a final concentration of 1 nM for all negative controls. Cells were treated with siRNA in antibiotic free DMEM/F12 (11330-032; Gibco) supplemented with 10% fetal bovine serum (FBS; Phoenix Scientific; PS-100) for 8 hours. siRNA was washed out and replaced with DMEM/F12 (11330-032; Gibco) supplemented with 10% fetal bovine serum (FBS; Phoenix Scientific; PS-100) and 1% penicillin/streptomycin for 8 hours prior to serum depletion for 24 hours and subsequent SAG (200nM; SML1314; Sigma) treatment for 24 hours (48 hours total of serum depletion) before fixation and immunostaining.

### Generation of SMO:GFP D21 and T21 cell lines

A construct consisting of human Smo with a C-terminal GFP tag downstream of a CMV promoter on a pHUSH backbone (SMO:GFP) was created by Chris Westlake’s lab. HEK293T cells were transfected with the SMO:GFP construct and lentivirus packaging plasmids, psPAX2 and pMD2.G, using lipofectamine 2000 (11668-027; Invitrogen). HEK293T medium containing virus was collected and added to target cells (RPE-1 D21 and T21 cells) with 2µg/mL polybrene. Target cells were incubated with viral medium for 24 hours. After 24 hours, new viral medium was added to target cells and target cells were left to incubate for an additional 24 hours. Transduced cells were selected using 10µg/mL puromycin for 3 days. Cells were flow-sorted to isolate GFP-positive populations.

### Fluorescence recovery after photobleaching

Live imaging dishes were generated by cutting out an 18mm diameter circle from the bottom of a 35mm tissue culture dish. A 30mm glass coverslip was then affixed to the bottom of the 35mm tissue culture dish using aquarium sealant. Dishes were stored at room temperature for 24 hours to allow sealant to cure, then UV-sterilized for two hours. The glass-bottom well in the dish was coated with collagen (C9791; Sigma) and incubated overnight at 4°C. Cover slips were then washed three times PBS before plating cells. SMO:GFP-D21 and SMO:GFP-T21 cells were plated at 200,000 per dish in DMEM/F12 (11330-032; Gibco) supplemented with 10% fetal bovine serum (FBS; Phoenix Scientific; PS-100) and 1% penicillin/streptomycin. To induce ciliation, medium was replaced with serum-free FluoroBrite DMEM (A18967-01; Gibco) supplemented with 1% penicillin/streptomycin for 20 hours prior to treatment with 200nM SAG (SML1314; Sigma) for 4 hours (24 hours total in serum depletion with 4 hours of SAG). Dishes were imaged live using Nikon Eclipse Ti-E microscope equipped with a controlled environment stage (OkoLab; H301-K-Frame; 37°C; 5% CO_2_). For all FRAP experiments, a 488nm laser was used. For Figures 3C and 3E, photobleaching was achieved using 60% laser power over a duration of 6 ms using a single point that encompassed the indicated area. For Figure 3D, photobleaching was achieved using 30% laser power over a duration of 6 ms along a line centered in the indicated area. For all FRAP experiments, three images prior to and one image after photobleaching were taken every 1 s. During the recovery acquisition, images were taken every 10 s for a total duration of 6 minutes. Fluorescent intensity was quantified in indicated regions and normalized to minimum intensity (at time of photobleaching) and the maximum intensity (prior to photobleaching) within each acquisition.

### Fluorescence microscopy

For FRAP, widefield images were acquired with a Nikon Eclipse Ti-E microscope (Nikon) equipped with a 100x Plan Apochromat objective (NA 1.40) and a Teledyne Photometrics Kinetix 22 sCMOS camera (Teledyne Photometrics). Confocal images were acquired with a Nikon Eclipse Ti inverted microscope stand equipped with a 60x Plan Apochromat objective (NA 1.40), Teledyne Photometrics Prime95B sCMOS camera (Teledyne Photometrics), and CSU-X1 (Yokogawa) spinning disk. Slidebook 6 digital microscopy software was used for confocal and widefield image acquisition. All confocal images were acquired at room temperature. All widefield images were acquired at 37°C and 5% CO_2_ for live imaging acquisitions. Identical acquisition settings were used for quantitatively compared images.

### Fluorescence quantification

To quantify the percent of cells with a primary cilium, cilia were labeled with ARL13B primary antibodies. Cilia ROIs were generated by particle analysis of objects resultant from thresholding ARL13B signal. PCNT intensity was quantified using a 5µm radius ROI centered on a CEP192 centroid. SMO intensity at the centrosome was quantified using a 2.5µm diameter ROI centered on a CEP192 centroid. All fluorescence intensity analyses were performed by recording the mean grey value of defined ROIs in ImageJ using maximum projected z-stacks. Due to variability, in-cell fluorescence intensity background subtraction was performed by subtracting the average intensity of three regions within the cell from the intensity in the defined ROI of the same cell. Fluorescence intensities were normalized for each cell to the average value of the untreated control condition in each respective experiment (i.e. D21; 24-hour serum depletion; no SAG). For Figure S2G, SMO intensity was measured along the cilium and PCNT intensity was measured in a 5µm radius centered on the centrosome of the same cell. A Pearson correlation was calculated for the paired PCNT and SMO intensities (n=3; 35 cells/replicate). For image averaging in Figure 3F and Figure 4F, 2.5µm radius image crops centered on CEP192 centroids were oriented to align cilia. Pixel values of SMO signal in each image crop were normalized. Normalized image crops were concatenated into a single stack per treatment group, and the average intensity was projected through the stack. SMO localization in peripheral regions was quantified (Figure 3H and 4G) according to the schematic in Figure 3H. Values derived from “Peripheral Signal” regions were averaged for each cilium and normalized to those of the D21, no SAG condition. For Figures 3G and S3C, line scans were performed along the cilia structure (2.5µm lengths from centrosome centroid along cilium) of the image averaging stack. Intensity values from each line scan were normalized to the minimum and maximum within each image averaging stack. For Figure S3F, line scans along approximate cilia structures were used to quantify CEP192, IFT88, and SMO fluorescent intensity. Intensity values were normalized to the peak intensity of each channel along the line scan and binned based on relative position along cilium.

### Statistics

Data were organized with Microsoft Excel (Microsoft) and plotted and analyzed using Graphpad Prism (GraphPad). Data center values (outlined in black) represent averages of each biological replicate while error bars represent SD. For FRAP analyses, the EasyFRAP webtool (https://easyfrap.vmnet.upatras.gr (Koulouras et al., 2018)) was used to calculate T_1/2_ values for SMO:GFP-D21 and SMO:GFP-T21 at each region. T_1/2_ values were calculated from a single exponential term equation that was fit to experimental data. All experiments utilized at least three biological replicates. A Student’s two-tailed unpaired *t* test was used to test significance between two normal unpaired distributions. A One-Way ANOVA was used to test significance between 3 or more normal unpaired distributions. A Two-Way ANOVA was used to test significance between 3 or more normal unpaired distributions when two variables were present. Differences between distributions were considered statistically significant if *p* values were less than 0.05. *p* values, sample sizes, and statistical tests used for samples are denoted in respective figure legends.

## Data Availability

No large omics datasets or computer code are associated with this work. Raw imaging files are available upon request from the corresponding author.

## SUPPLEMENTAL FIGURE LEGEND

**Figure S1:**
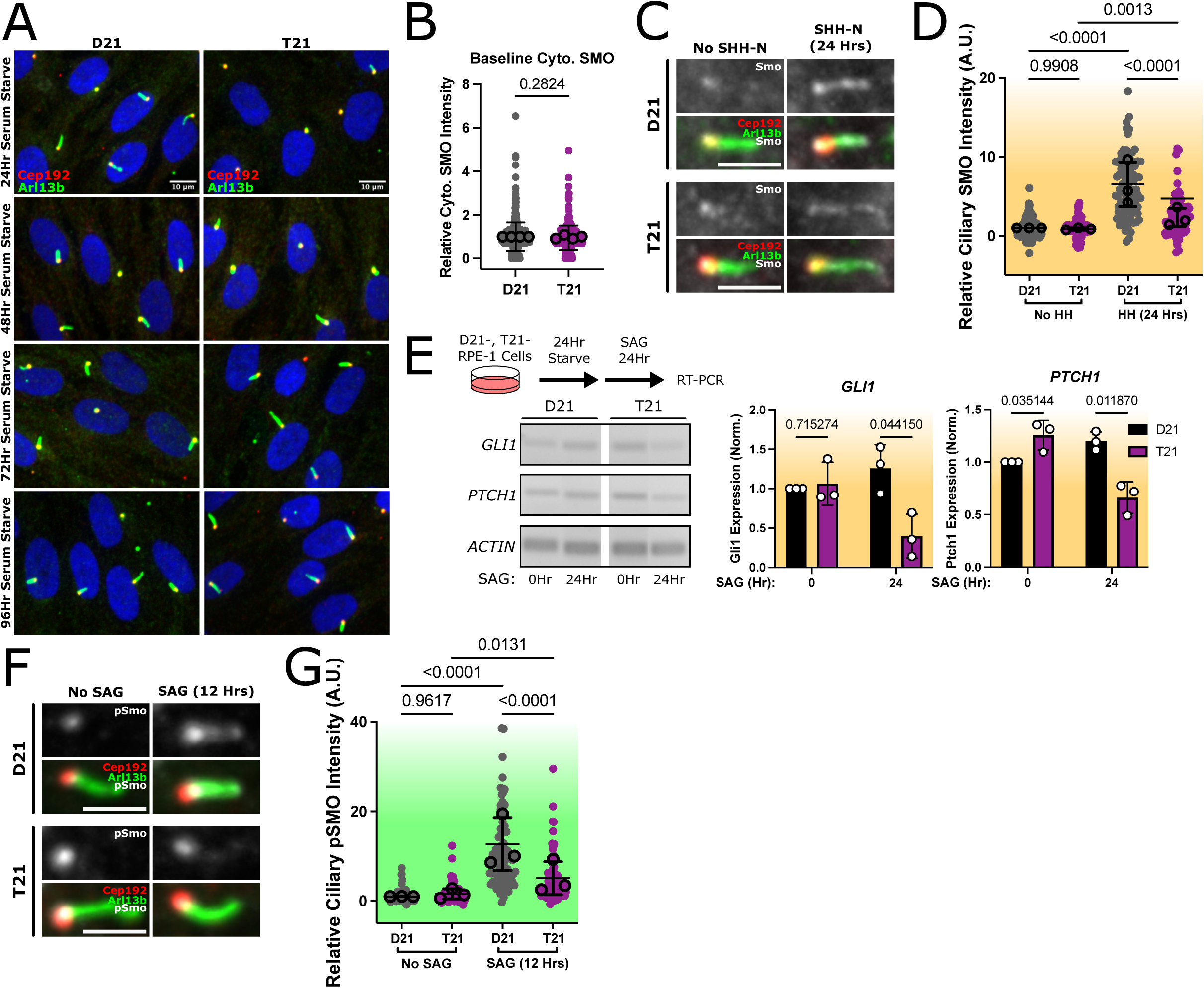
Trisomy 21 delays ciliogenesis and disrupts SHH transcriptional output. **(A)** Ciliation frequency is delayed in T21. Representative confocal images of ciliation frequency in D21 and T21 cells at various times of serum depletion. Cilia (ARL13b), green; centrosomes (CEP192), red. Scale bar, 10µm. **(B)** Baseline SMO levels are unchanged in T21 cells. Quantification of cytoplasmic SMO in D21 and T21 after 24 hours of serum depletion without SAG treatment (n=3; 30-150 cilia/replicate). **(C)** T21 decreases ciliary SMO levels compared to D21 cells after treatment with SHH-N. Representative confocal images of ciliary SMO in D21 and T21 cells at 24 hours post SHH-N treatment. Cilia (ARL13b), green; centrosomes (CEP192), red; SMO, grayscale. Scale bar, 5µm. **(D)** Quantification of ciliary SMO intensity in D21 and T21 post SHH-N treatment (24 hours; 48 hours total in serum depletion: n=3; 30 cilia/replicate). **(E)** T21 cells have a blunted transcriptional response to SHH stimulation. Experimental design to determine SHH transcriptional differences in T21 versus D21. Isogenic RPE-1 D21 and T21 cell lines were cultured in serum depleted medium for 48 hours with or without SAG (24 hours) before harvesting RNA for RT-PCR. Representative images of PCR amplification of *GLI1*, *PTCH1*, and *ACTIN* and quantification of *GLI1* and *PTCH1* PCR amplification with and without SAG treatment (n=3). P-Value, *t*-test. **(F)** T21 decreases ciliary pSMO levels compared to D21 cells. Representative confocal images of ciliary pSMO in D21 and T21 cells after 12hrs of SAG treatment (36 hours total serum depletion). Cilia (ARL13b), green; centrosomes (CEP192), red; pSMO, grayscale. Scale bar, 5µm. **(G)** Quantification of ciliary pSMO intensity post SAG treatment (12 hours; 36 hours total in serum depletion; n=3; 30 cilia/replicate). P-value, 2-Way ANOVA.

**Figure S2:**
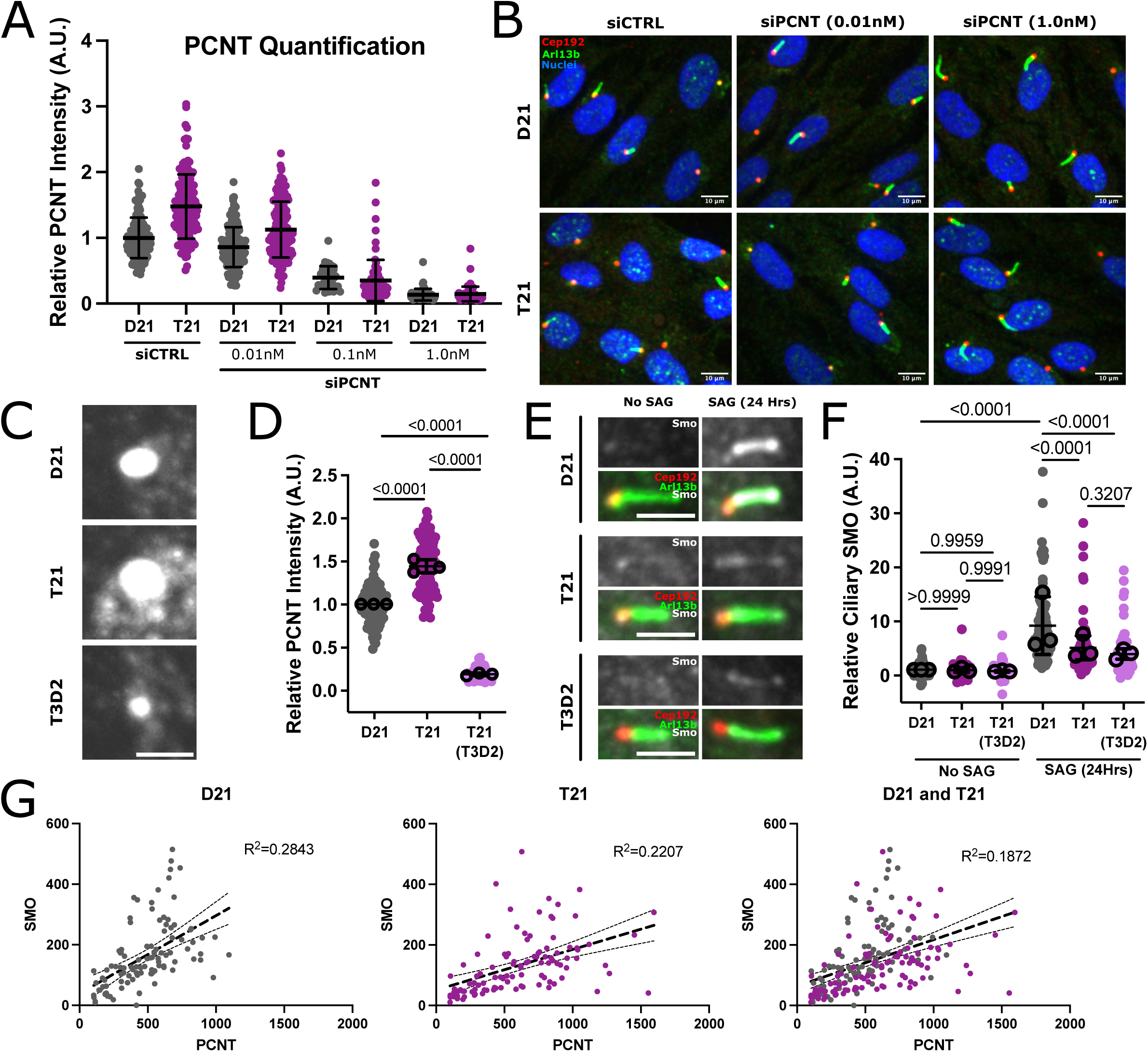
Elevated PCNT in T21 cells does not diminish ciliary SMO. **(A)** Titration of siPCNT to determine concentrations that reduce PCNT to approximately disomic and subdisomic levels. Quantification of PCNT levels at the centrosome in control and siPCNT-treated D21 and T21 cells. **(B)** Representative confocal images of ciliation frequency in control and siPCNT-treated D21 and T21 cells at 24 hours of serum depletion without SAG treatment. **(C)** Representative confocal images of PCNT at the centrosome in D21 and T21-T3D2 (2n PCNT) cells at 48 hours of serum depletion without SAG treatment. **(D)** Quantification of PCNT levels at the centrosome in D21 and T21-T3D2 (2n PCNT) cells (n=3; 30 cells/replicate). **(E)** Representative confocal images of ciliary SMO in D21 and T21-T3D2 (2n PCNT) cells after 24 hours of SAG treatment (48 hours total in serum depletion). Cilia (ARL13b), green; centrosomes (CEP192), red; SMO, grayscale. Scale bar, 5µm. **(F)** Reduction of *PCNT* to genomically disomic levels does not rescue ciliary SMO localization in T21 cells. Quantification of ciliary SMO intensity in control and D21 and T21-T3D2 (2n PCNT) cells with and without SAG treatment (n=3; 30 cilia/replicate). **(G)** Ciliary SMO levels are positively correlated with PCNT levels. Pearson correlation between centrosomal/pericentrosomal PCNT and ciliary SMO in D21 and T21 cells (n=3; 35 cells/replicate). P-value, 2-Way ANOVA.

**Figure S3:**
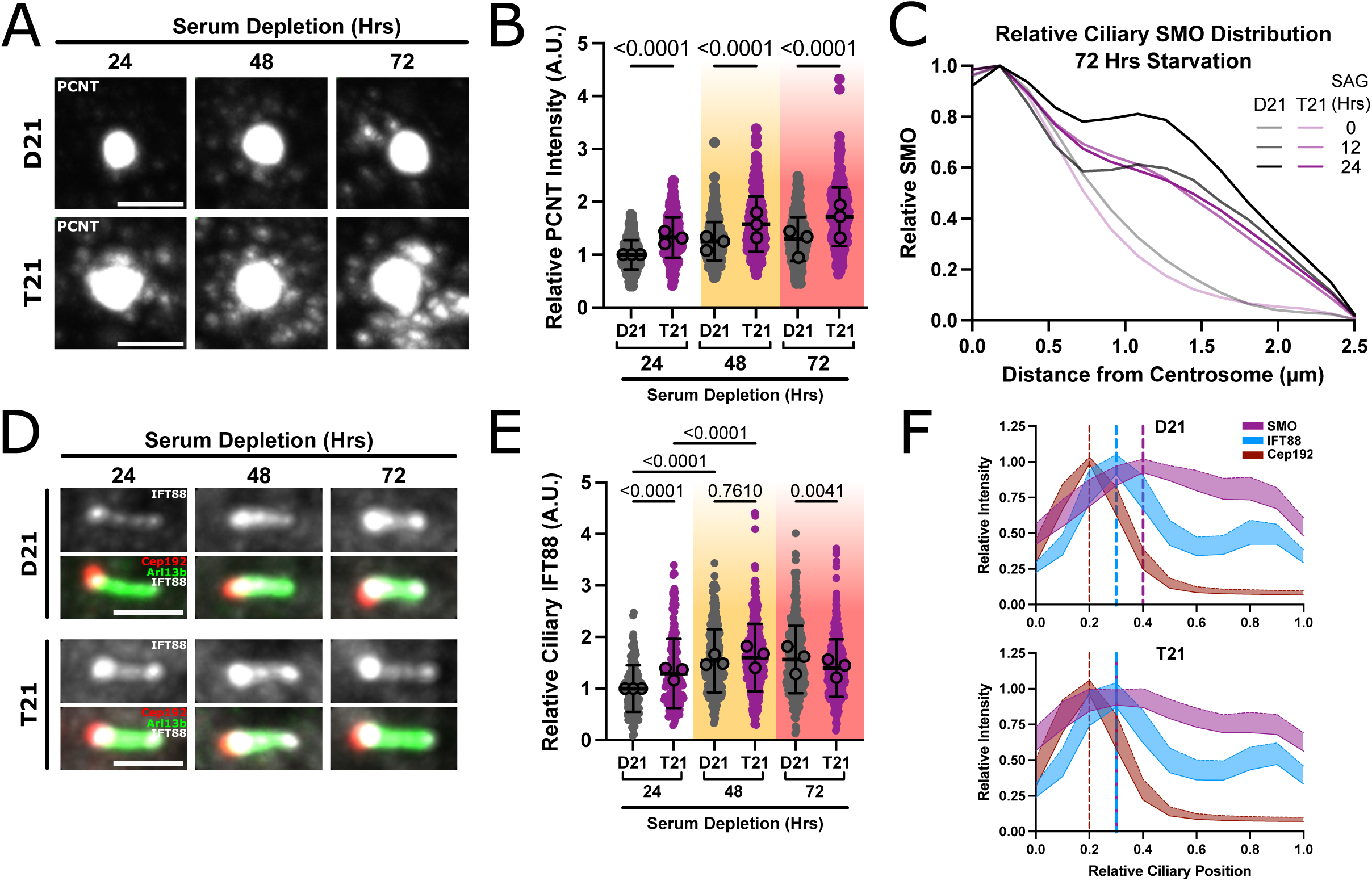
Localization of ciliary SMO in T21 increases with cilia maturation independent of PCNT. **(A)** PCNT levels at the centrosome do not change with increased serum depletion time. Representative confocal images of PCNT at the centrosome of D21 and T21 cells at various serum depletion times. Scale bar, 5µm. **(B)** Quantification of PCNT levels at the centrosome at various serum depletion times. Color gradients indicate serum depletion times (24 hours, no color; 48 hours, yellow; 72 hours, red; n=3; 50-120 cells/replicate). **(C)** Increased ciliary maturation time partially rescues SMO localization differences in T21 to D21 levels. Line scan analysis of SMO distribution along mature cilia. **(D)** T21 increases ciliary IFT88 but is rescued to D21 levels with increased cilia maturation. Representative confocal images showing ciliary IFT88 in D21 and T21 cells at indicated serum depletion times. Cilia (ARL13b), green; centrosomes (CEP192), red; IFT88, grayscale. Scale bar, 5µm. **(E)** Quantification of ciliary IFT88 in D21 and T21 cells. Color gradients indicate serum depletion times (24 hours, no color; 48 hours, yellow; 72 hours, red; n=3; 40-140 cilia/replicate). **(F)** Ciliary IFT88 distribution is unchanged while ciliary SMO distribution is restrained to the proximal end of cilia in T21. Normalized line scan intensities of IFT88 (blue), SMO (purple), and CEP192 (red). Normalized line scans were plotted along binned ciliary lengths (10% increments) to illustrate relative distribution along cilium. Dashed vertical lines indicate peak relative intensity (IFT88, blue; SMO, purple; red, CEP192). P-value, 2-Way ANOVA.

## ACKNOWLEDGEMENTS

We thank Andrew Lane and David Pellman for generously providing Hsa21 RPE-1 cell lines; Chris Westlake for the SMO:GFP construct; Andrew Holland for the CEP192 antibody; and Erik Collet for ImageJ Plugins used in Fluorescence Quantification. We acknowledge the Pearson lab for helpful discussions. C.G.P. is a member of the Linda Crnic Institute for Down syndrome. This study was supported in part by the National Institutes of Health P30CA046934 funded Flow Cytometry Shared Resource (RRID:SCR_022035). This research was funded by R35 GM140813 to C.G.P., the NSF Graduate Research Fellowship Award (DGE-2439026) to W.E.S., the Damon Runyon Cancer Research Foundation (Merck Fellow DRG-2478-22) to C.E.J., and the CU Anschutz Molecular Biology Bolie Fellowship Award to B.L.M.

## AUTHOR CONTRIBUTIONS

Conceptualization: C.G.P., W.E.S.; Methodology: W.E.S., B.L.M., C.E.J.; Investigation: W.E.S., B.L.M.; Formal Analysis: W.E.S., B.L.M.; Data Curation: W.E.S., B.L.M.; Resources: C.G.P., C.E.J.; Writing – Original Draft Preparation: W.E.S.; Supervision: C.G.P.; Funding Acquisition: W.E.S., C.G.P.

## Abbreviations

(T21): Trisomy 21
(SHH): Sonic Hedgehog
(IFT): Intraflagellar Transport
(GPCR): G-protein Coupled Receptor
(SMO): Smoothened
(PTCH): Patched
(PCNT): Pericentrin

